# Early-life experience determines the social stability of adult communities in female mice

**DOI:** 10.1101/2023.11.29.569157

**Authors:** Birte Doludda, Jadna Bogado Lopes, Sara Zocher, Gerd Kempermann, Rupert W Overall

**Author notes:** Corresponding author. (RWO). These authors contributed equally to this work (Birte Doludda, Jadna Bogado Lopes). These authors contributed equally to this work (Gerd Kempermann, Rupert W Overall).

## Abstract

Early social interactions critically shape lifelong individual behaviour but their development and stability have been difficult to study in laboratory settings. Our experimental platform allows the automated real-time and longitudinal study of social structure in mice living in a shared environment, providing previously inaccessible behavioural measures. These include estimates of directed agonistic activity for the reconstruction of social hierarchies and their dynamics over the life-course. In an all-female colony of genetically identical mice, pairwise inter-individual interactions revealed stable dominance hierarchies that already emerged early in life. Older animals introduced into a new colony exhibited a steeper hierarchy compared to adolescent mice or mature mice that had been co-housed throughout life. Our longitudinal analysis of social dominance hierarchies highlights the critical role of early-life experience.

**One-Sentence Summary:** Automated tracking of mouse colonies revealed differences in social structure dependent on the animals’ early-life experience.

## Main Text

The environment to which an animal is exposed in early life exerts a defining influence on its behaviour as an adult (*1, 2*). Current theory distinguishes the shared external environment from the individual’s response to that environment; the non-shared component. This non-shared experience consists of exposure to different environments as well as individualised exploration of shared spaces (*3*–*7*). The role of social influences in shaping individuality, however, has been less well studied—despite an acknowledgement that early-life social interactions influence behaviour and health in adulthood (*8*–*10*). Whereas genetics and the physical environment can be readily controlled, all remaining individual phenotypic variation has often been considered stochastic effects (*11*). In contrast, we propose that social experience is a critical factor that can be dissociated from other components of the non-shared environment. We have developed an experimental platform that exposes the non-shared environment by controlling both genetic background and shared environment—allowing diverging individual behavioural trajectories to emerge (*5, 6, 12*). In this system, the free exploration of individual mice in a large network of cages is tracked with high resolution over extended periods of time. We have now extended this platform to study the development of individual social behaviour and social hierarchies. The ability to follow pairwise (dyadic) inter-individual interactions at high resolution and throughout life in an experimental population will open up a new paradigm in the laboratory study of social development.

Most existing techniques used to study social interaction under laboratory conditions rely on specialised observation arenas, which constitute an unnatural environment that can significantly alter behaviour (*13, 14*). Home cage monitoring has been successful in allowing social groups of rodents to be followed without disturbance from researchers (*15*–*18*), although the size of the colonies is still restricted. Observation periods in those settings also tend to be relatively short, hampering the investigation of social hierarchies—which are established through repeated social interactions over longer time periods rather than isolated social encounters (*19, 20*). Animals in a large colony do not necessarily all interact with each other and subordinate members may avoid high-ranking individuals to spare the unnecessary expenditure of energy in a contest they are assured to lose (*21*). Our paradigm allows individual mice in large colonies to be tracked with high spatial and temporal resolution in their home cage environment and over long periods (*6, 12*). Our new analysis pipeline reveals a wide range of previously inaccessible behavioural measures, including estimates of directed agonistic activity that allow the reconstruction of social hierarchies and their dynamics.

We developed an open-source software package, ‘ColonyTrack’, to facilitate data pre-processing and feature extraction (**fig. 1A**). This software identifies the location of every animal at every point in time (**fig. 1B**), from which measures of social interaction can be extracted. Actograms (**fig. 1C**) show any activity anomalies and missing data. The calculation of metrics (derived features) is based around three core behaviour types: ‘activity’ (including distance moved, speed, number and times of peak activity), ‘exploration’ (the distribution of time spent in each cage) and ‘sociality’ (measures of time spent with other animals). An automated ethogram shows a scaled representative metric from each of these three classes calculated hourly and presented in a 3-component subtractive colour model to provide an intuitive assessment of the experiment (**fig. 1D**).

**Fig. 1.**
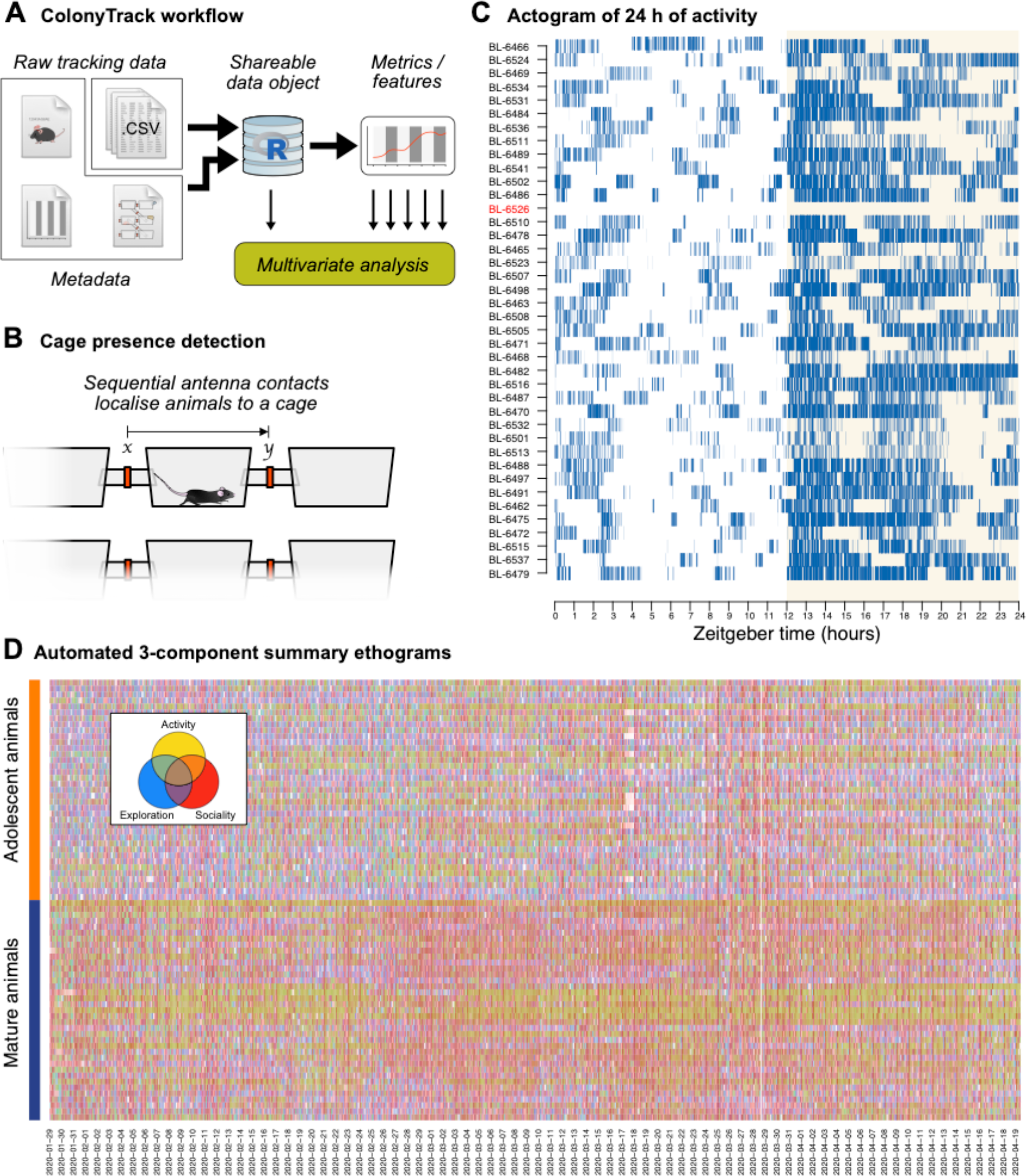
The ColonyTrack software extracts multiple features from longitudinal home cage monitoring. (**A**) Data analysis workflow. Raw data and metadata are efficiently and reproducibly processed into a shareable data archive. Subsequent feature extraction yields 21 metrics that enable rich multivariate analyses. (**B**) Cage presence detection of individual animals allows estimation of social interactions within the colony. (**C**) An actogram for the adolescent colony covering one 24-hour period. One inactive (missing) animal has been automatically flagged with a red *y* axis label. The x axis is in Zeitgeber time (ZT) starting at lights-on with the dark (active) period shaded in pale yellow. (**D**) An automated ethogram provides a rapid overview of the 3-month-long experiment. Three colour channels representing the three core behavioural axes (yellow: activity, red: social interaction, blue: exploration; see inset) are mixed to provide an intuitive overview of the total behaviour across very many animals with a dense 1-hourly resolution. The pronounced yellow component in the mature colony, for example, draws attention to increased activity in these animals, especially in a subset of individuals.

Calculation of the social hierarchy was done using measures of dyadic agonistic interactions. These were estimated from cases where two individuals ran after each other through the same cage (**fig. 2A**). We had noted that a proportion of such ‘following events’ were agonistic, with the non-agonistic background being evenly distributed and thus normalised out during hierarchy calculation (**fig. 2B**; see Online Methods for details). To confirm the contribution of agonistic behaviour to the ‘following’ counts, we performed an initial experiment with a highly diverse population (outbred CD1 mice). Manual ethology of over 170 hours of video (**fig. 2C-D**) established that the observed interactions were either neutral (the non-agonistic background that occurs between all animals regardless of hierarchy position) or agonistic (including biting, brief fights or defensive posturing), with very few cases of agonistic behaviour in the reverse direction (where the followed animal was the aggressor). A hierarchy built from these agonistic data (forward and reverse) closely mirrored the hierarchy calculated by the ColonyTrack software (**fig. 2E**). We selected a subset of individuals (5 highest-ranking and 5 lowest-ranking, as calculated by ColonyTrack) and performed a traditional out-of-cage tube test at the end of the experiment. Excepting one animal (the lowest-ranked individual, which won all of its trials), these dominance scores matched those from ColonyTrack (**fig. 2F-G**). These results confirm that a working estimate of social hierarchy can be obtained from fully-automated home cage tracking.

**Fig. 2.**
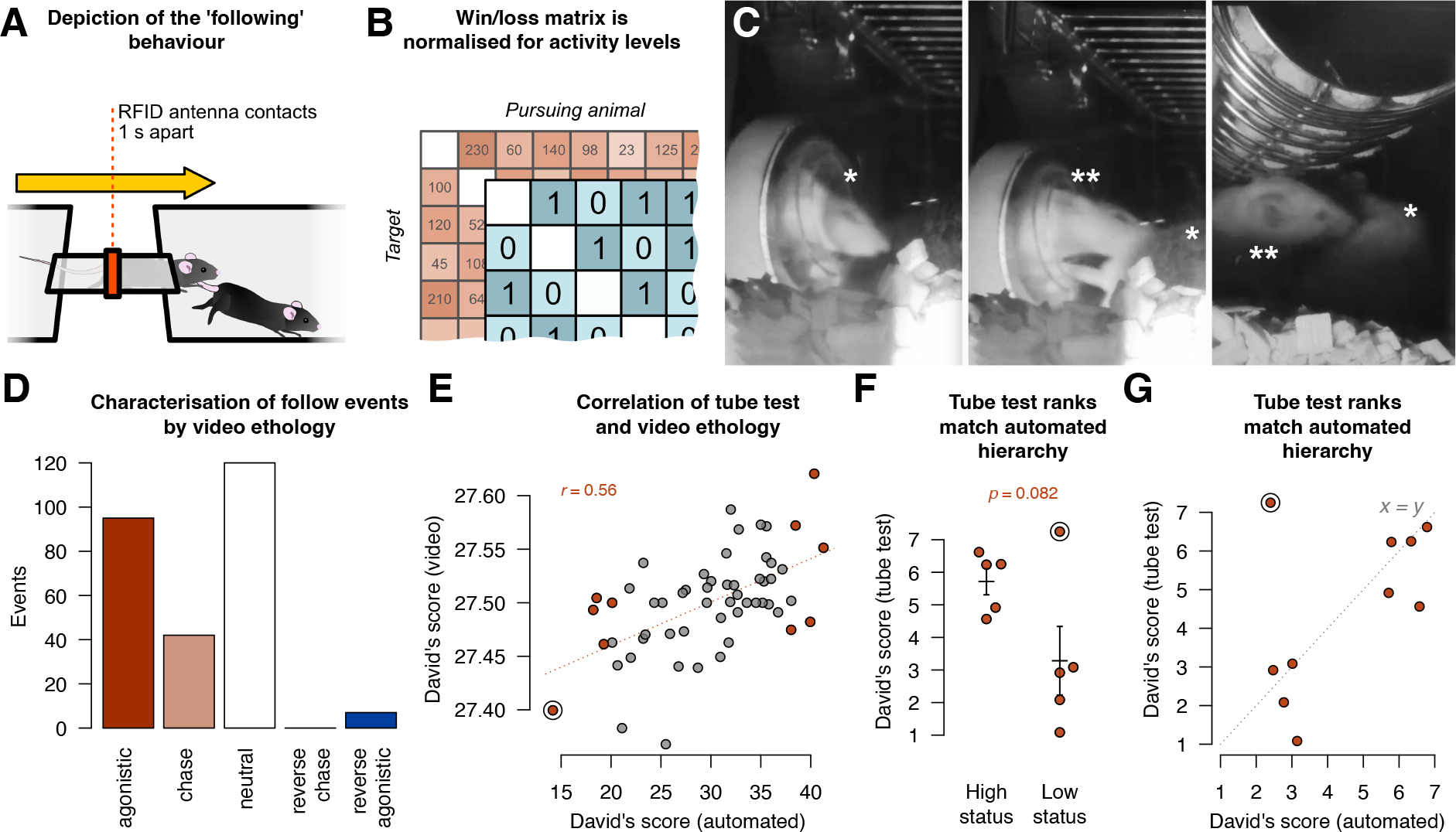
Social dominance can be reliably estimated from automated home cage tracking. (**A**) Following events are when two animals pass in the same direction within one second of each other. (**B**) Although many such interactions also occur by chance, there is a bias such that dominant animals in a dyad are more often the follower. This means that when raw following data (red grid) are discretised into a ‘win’ or ‘loss’ for each day (blue grid), the background activity levels are normalised out. (**C**) A typical agonistic interaction can be observed in video footage where the target animal (frame 1; asterisk) is pursued at speed by the aggressor (frame 2, double asterisk) often leading to clear agonistic behaviour, such as a fight (frame 3). (**D**) Video footage of all ‘follow’ interactions for one cage were classified manually. Alongside the background ‘neutral’ interactions (white bar), most were indeed agonistic (red). A detailed discussion can be found in the Supplementary Information. (**E**) The video ethology-based hierarchy correlated well with the automated estimates. The focus animals used for the tube test are highlighted in red. (**F**) The tube test clearly showed the highest- and lowest-status animals (from automated estimates) to be correspondingly ranked—excepting one animal, the most subordinate animal in the home cage, which behaved unusually aggressively in the unfamiliar setting of the tube test. This individual is shown with a black ring in panels E–F. See Supplementary Information for a comment on problems with traditional out-of-cage testing. (**G**) The dominance ranks for the 10 focus animals from the tube test matched those from the ColonyRack (the *x = y* line is shown for comparison).

To address the role of the social environment in early life individualisation, we next studied two colonies of female mice in our system. In one colony (‘adolescent’), the animals were introduced at 5 weeks of age and maintained in the home cage system throughout their adolescence. In the second colony (‘mature’), animals were already 6 months old before being co-housed. All animals were tracked continuously for three months. We also applied our new software to reanalyse data from an earlier study (*6*) in which mice were tracked from weaning over an age range that overlapped both colonies of the current experiment.

The 21 metrics delivered by ColonyTrack provided a longitudinal multi-parametric behavioural profile describing how the animals responded to their physical and social environment (**fig. S1**). As we saw in the preliminary experiment, we noted that higher activity resulted in more chance following encounters (confirmed by randomizing the actogram data; **fig. S2**). Because following behaviour is directed, all encounters can be summarized in a win/loss table and enables animals to be ranked using the Davids’ score (*22, 23*). The distribution of ranks for each animal in the adolescent colony over all days showed that the most dominant animals exhibited very different average ranks than the least dominant (**fig. 3A**) and this trend was even more pronounced in the mature colony (**fig. 3B**). This was in contrast to the established colony (**fig. 3C**) and randomised data where ranks were broadly distributed around the average value (**fig. S3**). Whereas there were only a few animals in the adolescent colony that stood out as dominant, there were 11 animals for whom the lower quartile of the dominance score was never less than the mean value for the whole colony (red lines in **fig. 3A, B**; **fig. S3**). We labelled these 11 most dominant animals the *alpha* group and found that their (social) behaviour stood out from the rest of the colony.

**Fig. 3.**
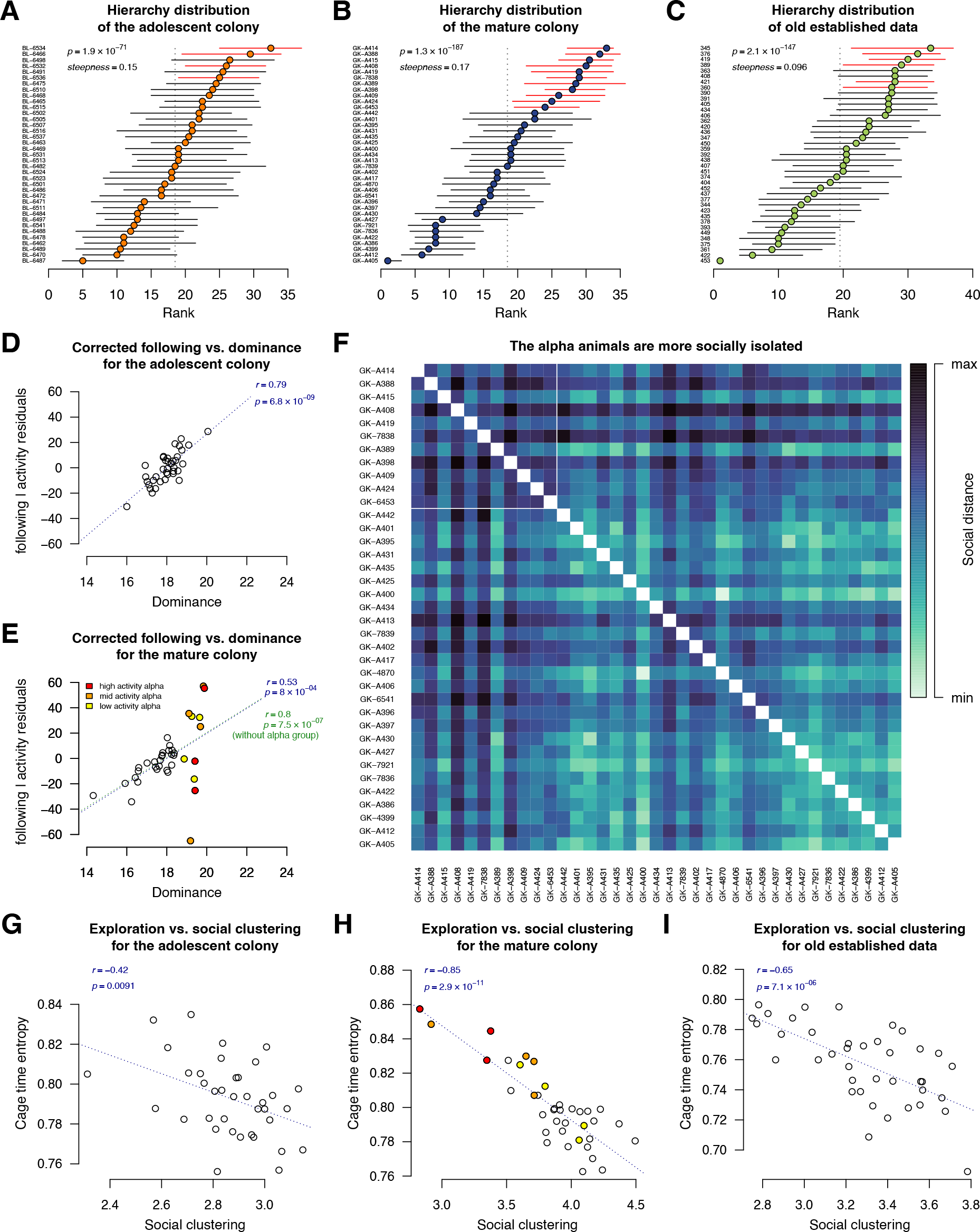
A re-established colony is characterised by a top-heavy hierarchy. (**A–C**) The hierarchy rank distribution of each animal is shown as median ± quartiles over all days. Animals in each plot are ordered by their median rank over all days. The average rank for the colony is shown as a dotted vertical line. Colours match the colony colour codes in fig. 2. (**D, E**) ‘Following’, after correction for background activity levels (see fig. S2), is predictive with dominance for all animals except the *alpha* group. Regardless of activity, the *alpha* group individuals are at the top of the social hierarchy. Regression line and *r*^2^ in green in panel E exclude the *alpha* animals. (**F**) A pairwise clustering matrix shows that the *alpha* individuals are typically more socially distanced from other animals. Rows and columns are ordered by dominance score and thin white lines delineate the *alpha* group in the top left corner. (**G–I**) Exploratory behaviour (cage time entropy) inversely correlates with social clustering. In panels A-C, *p*-values from Friedman tests and the steepness scores are shown, in all other panels, *p* and *r* are from Pearson correlations.

Plotting the activity-corrected following metric against the David’s dominance score for all three colonies revealed a consistent linear relationship for all animals with the exception of the *alpha* group. Whereas in the adolescent colony, the position in the hierarchy was clearly related to the amount of agonistic behaviour (**fig. 3D**), in the mature colony, the *alpha* individuals exhibited a wide range of activity levels (**fig. 3E**). This indicated that the most dominant animals in the mature group were characterised by high inter-individual behavioural variability.

We further asked whether this dominant *alpha* group was a coherent social cluster that predominantly interacted with each other. Social distance was calculated as the shortest path (in terms of cage units) between two animals at any time. We discovered that the *alpha* animals were in fact among the most socially distanced individuals, both from the bulk of the colony as well as from each other (**fig. 3F**). Using ethograms taken by a human observer, we had previously reported an inverse relationship between social and environmental exploration (*24*). Although in the present study we did not find such a correlation in the adolescent mice, the highly explorative animals in the mature colony spent less time with other mice (social clustering) and showed a higher average distance to other mice (social distance)—both indicative of reduced sociality (**fig. 3H–I**; **fig. S4**). Together, these data suggest that in female mice social dominance is associated with a reduction in social contacts and interaction.

To determine whether the social hierarchy developed during the time of co-housing or was determined already at the start of the experiment, we compared the change in social dominance scores between the first and last week of the experiment at the individual mouse level. In the adolescent colony, the change in social rank was distributed evenly throughout the different positions in the hierarchy (**fig. 4A**), whereas in the mature colony there was a tendency for higher-ranked individuals, including the *alpha* group, to further gain in rank and for lower-ranked individuals to lose rank (**fig. 4B**). These results show that the mature colony not only remained more stable in terms of ranking but that the hierarchy also became more polarised over time. Correlation of the David’s dominance scores from the first and last week corroborated this finding, with a poor correlation (*r*^*2*^ = 0.1) in the adolescent colony (**fig. 4C**) but a stronger relationship (*r*^*2*^ = 0.3) among the mature animals (**fig. 4D**). These results suggest that, unlike animals raised throughout life together, female mice grouped in later life rapidly establish a hierarchy which becomes even stronger during further co-housing.

**Fig. 4.**
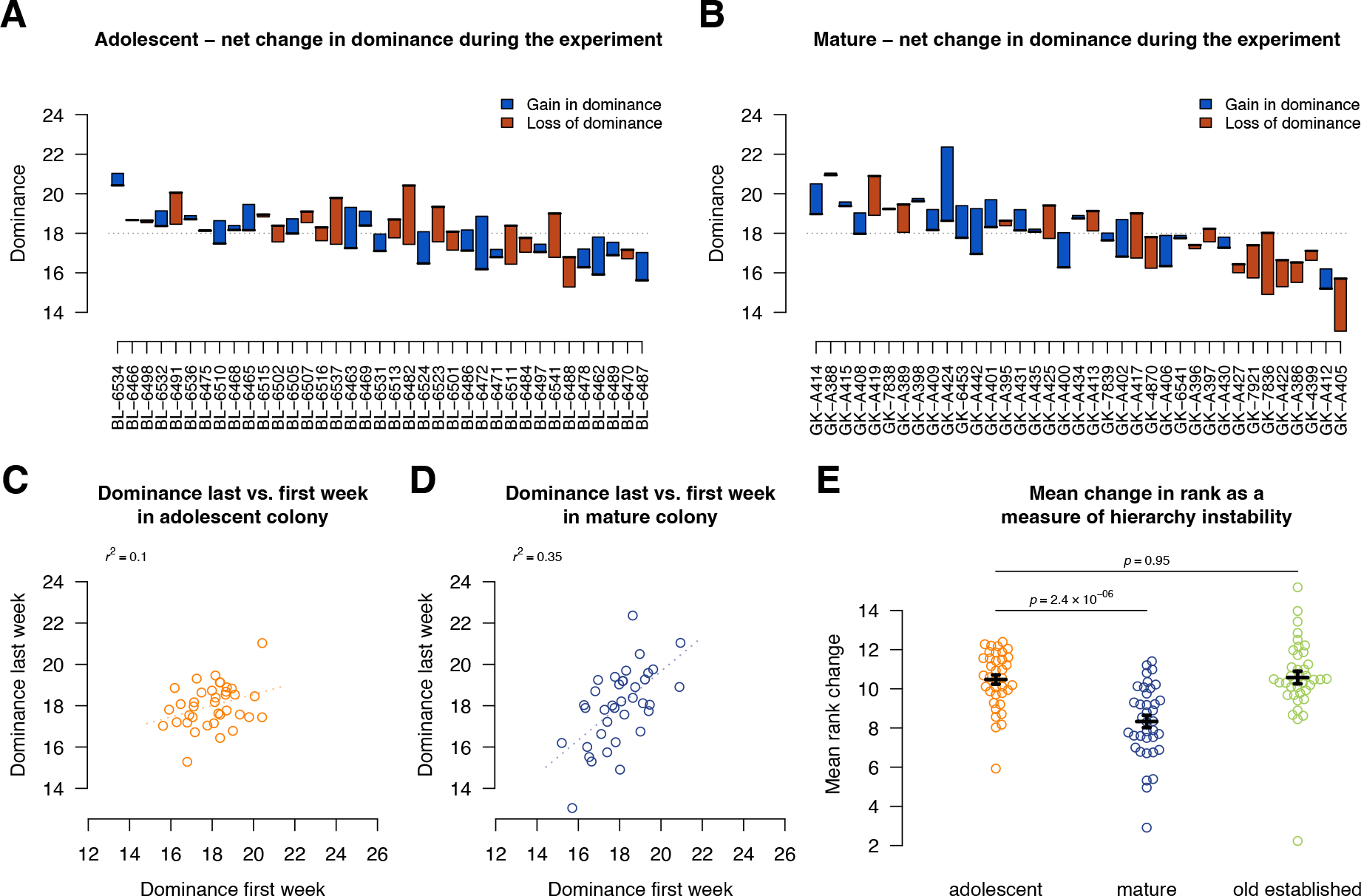
A re-established social hierarchy is more stable and more polarised. The dominance (David’s scores) were calculated over the first and last weeks of the experiment for both the adolescent (**A**) and mature (**B**) colonies and plotted as differences—blue bars show an increase and red bars show a decrease in dominance over time. Animals are ordered by mean dominance throughout the whole experiment (as in fig. 3). Last -vs. first-week dominance scores for the adolescent colony were not correlated (**C**) unlike the mature colony (**D**). When the absolute change in rank at consecutive timepoints was summarised (mean over all days), the animals of the mature colony exhibited less hierarchy instability than either the adolescent or a similarly-aged established colony (**E**). Reported *p* values are from the Dunnett test.

As a measure of rank stability over the entire experiment period, we calculated the mean absolute rank difference for each animal (see Online Methods for details). The mature animals had generally less change in rank compared to the adolescent animals (mean of 8.3 vs. 10.5; *p* = 2.4 × 10^−6^; Dunnett test; **fig. 4E**), again indicating that the mature hierarchy was more stable. A comparison with data from the last 70 days of the established colony from our previous study (corresponding to the period covered by the mature dataset) indicated that those animals, that were of an equivalent age but had been together as a colony since weaning, were essentially indistinguishable from the adolescent colony (mean of 10.6 vs. 10.5; *p* = 0.95; Dunnett test; **fig. 4E**). This last finding was intriguing as it suggests that the difference in the mature colony was not due to age or length of exposure to the ColonyRack environment but was rather a result of the animals’ social environment during early life—the only factor characterizing both the adolescent colony and the established colony from our previous study (*6, 24*). These results argue for the importance of early-life social structure on the development of individual behavioural trajectories in female mice.

There is evidence that environmental enrichment in early life can lead to reduced dominance in rats (*25*). The complexity of the ColonyRack system represents an enriched environment, which may have suppressed dominant behaviour and hierarchy building in the early-established colonies as has been reported in the more constrained standard laboratory housing conditions. It should be noted, however, that all animals in this study spent at least 3 months in the ColonyRack, showing that enrichment in late life was not sufficient to acutely alter social dominance behaviour. An alternative hypothesis is that the hierarchies prior to our experiment were not purely linear in structure, but dominated by a few high-status individuals in what is known as a ‘despotic’ hierarchy (*26*). If such a social structure arises during early-life de-novo hierarchy development, then a flat, unstable hierarchy with very few high-ranked outliers would be expected. However, by sampling many small standard-housed colonies, as we did to create the late-life mature colony, multiple ‘despots’ would likely be brought together, who would encounter discord between their perceived rank (established in their previous environment) with their position in the new hierarchy. Previously highly-ranked animals finding themselves further down the hierarchy may be motivated to increase their agonistic behaviour to re-establish their perceived rightful position, while animals retaining a high rank might retain that rank without conflict. This pattern is consistent with the wide range of agonistic behaviour levels observed in the *alpha* group in this study, but needs further work to confirm.

Using the experimental paradigm presented above, the present work proposes a role for social interaction in the development of animal personality and provides valuable background for the design of further research in the field of individuality. The open availability of our analysis software will enable this previously understudied aspect to be further explored by the community in future studies. We have shown that female mice brought together as adults established a distinct and stable hierarchy that was not present in animals co-housed throughout life.

Furthermore, the animals higher in the social hierarchy tended to be more socially isolated and more explorative than the remainder of the colony. The stability of these distinct behavioural profiles has revealed a previously underexplored aspect of animal personality. These results highlight the crucial influence of the social environment during early life on social behaviour and individuality later in life.

## Acknowledgments

We thank Sandra Günther, Anne Karasinsky and the animal facility staff at the DZNE, Dresden for their dedication and care of the mice.

## Author contributions

Formal Analysis: RWO

Methodology: BD, JBL, SZ, GK, RWO

Investigation: BD, JBL, RWO

Funding acquisition: GK

Supervision: GK, RWO

Writing – original draft: RWO

Writing – review & editing: BD, JBL, SZ, GK, RWO

## Data and materials availability

The ColonyTrack R package is available at: https://github.com/rupertoverall/ColonyTrack/; data are available at: https://doi.org/10.5281/zenodo.8333908

## Supplementary Materials

Materials and Methods

Figs. S1 to S4

Supplementary Text

Data S1 to S9

