## Supplementary Materials for "Early-life experience determines the social stability of adult communities in female mice"

**This PDF file includes:**

Materials and Methods
Figs. S1 to S4
Supplementary Text

**Other Supplementary Materials for this manuscript include the following:**

Data S1to S6 Raw data and metrics from the *ColonyTrack* software.

### Materials and Methods

Animals and housing

The mice used in the preliminary experiment were 60 female outbred CD1 mice purchased from Janvier Labs, which were 4 weeks old on arrival in the laboratory. Each animal had a RFID microtransponder (SID 102/A/2; Euro I.D.) injected subcutaneously under brief isoflurane anaesthesia and one week later was introduced to a 70-cage ColonyRack system and co-housed and RFID-tracked for 3 months.

The mice used in the main study were females of the strain C57BL/6JRj purchased from Janvier Labs. These animals were either 4 weeks (adolescent colony) or 23 weeks (mature) old on arrival in the laboratory and were introduced to the home cage system one week later, after being tagged with a subcutaneous RFID microtransponder (SID 102/A/2; Euro I.D.) under brief isoflurane anaesthesia. Animals were housed, as in our previous study(*1*) in one half of a ColonyRack home cage system (PhenoSys, Berlin) such that each colony had access to 35 connected Euro Type-II cages connected by Plexiglas® tunnels fitted with RFID antennae. Mice were maintained on a 12:12 hour light:dark cycle with ad libitum access to food and water on each level of the cage system. About half of the cages were provided with 'toys' (plastic tubes and shelters) which were moved on a weekly basis to provide a degree of environmental enrichment. All experiments were conducted in accordance with the applicable European and national regulations (Tierschutzgesetz) and were approved by the local authority (Landesdirektion Sachsen; file number TVA 16/2018).

Tube test

At the end of the ColonyRack tracking period, 10 focus animals from the CD1 experiment were selected (the 5 highest-ranking and the 5 lowest-ranking individuals based on the David's score from the automated 'following' metrics) for out-of-cage assessment in the tube test. The single tube dominance apparatus consists of two transparent plastic boxes connected with a transparent Plexiglas® tube (diameter 30 mm, length 385 mm). A slit in the center of the tube allowed manual insertion/removal of a transparent barrier. Bedding material from the home cage was placed inside the two compartments to simulate their natural environment and reduce stress. Mice were removed on day one and placed randomly in one compartment and gently encouraged to walk through the tube. This was performed with each mouse twice, randomizing the site of entry, to habituate them to the environment. Over the following 5 days, each pair of animals (45 dyadic interactions for the 10 focus animals) was tested once per day. For each trial, the two mice were placed at each end according to a randomised schedule. When both mice entered the tube and reached the divide, it was pulled out and the tournament was started. The trial ended when one mouse (the loser) left the tube with all four paws. The resulting win/loss table for all 225 interactions was used to calculate the David's scores.

Video ethology

A camera (C920 Pro HS Webcam from Logitech) recorded one cage of the ColonyRack. In order to obtain 24-hour recording (including the dark periods), the infrared filter was removed and an external infrared light source was used to illuminate the cage of interest. Video footage was collected over the 5 days immediately prior to the tube test. Timestamps of follow events into and out of the focus cage were identified by the *ColonyTrack* software and were used to extract video clips of the event. Two researchers (BD and RWO) reviewed the video clips and scored each interaction as 'neutral' (no interaction between the two following animals), ‘agonistic’ (biting, fighting by the follower or clear defensive posture by the target animal), ‘chase’ (clear chasing behaviour), ‘reverse chase’ (chasing in the opposite direction as detected by the software—this should not, and indeed did not, occur) and ‘reverse agonistic’ (where the target animal turned on the chaser and initiated agonistic behaviour). Interactions that were not clearly visible (e.g. due to other animals obscuring the view) were discarded from this analysis. For analysis, the ‘chase’ category was not considered (not neutral, but also circularly defined due to the selection of follow events).

Statistical analysis

Preprocessing of the RFID tracking data was done using our custom-built software package, *ColonyTrack*. The analysis for the current study used *ColonyTrack* version 1.0.4. The *ColonyTrack* software is available from https://github.com/rupertoverall/ColonyTrack and a detailed description of all metrics as well as full documentation can be found at https://rupertoverall.net/ColonyTrack/. The metrics investigated in this study were: activity (called ‘distance moved’ in *ColonyTrack*), which is the number of times an animal moved from one cage to another; ‘cage time entropy’, a measure of explorative behaviour calculated from the Shannon entropy of the likelihood of an animal being found in a particular cage (this is related to the ‘roaming entropy’ used in other publications (***1*, *2***), but calculated using the total time in each cage rather than from discrete sampling of timestamps); ‘social clustering’, which is the average number of other animals in the same cage as the focus animal; ‘social distance’, which is the average number of cages separating the focus animal from its cage mates; ‘follow events’, defined as two animals passing through the same tunnel in the same direction within 1 s of each other; and ‘follow dominance’, which is the fraction of follow events where the focus animal was the follower versus where it was being followed. Except where explicitly noted, the more robust David’s score was used to report dominance values, see below.

All analysis was performed in R/BioConductor (***3***). As the raw metrics were not normally distributed, non-parametric statistics were used unless indicated. Standard deviations in figure 2 were calculated by first log-transforming the activity score ('distance.moved'; panel A) and then max-min standardisation (panels B, F and H). Normalised David's scores, with correction for chance, were calculated using the R package *steepness* (***4*, *5***). A score for hierarchy stability was derived by first calculating the rank order of David's scores for each day and then calculating the change in rank for each animal (the absolute value of the rank at time *t* minus the rank at time *t - 1*). The mean absolute rank change was taken as the hierarchy stability score for that animal.

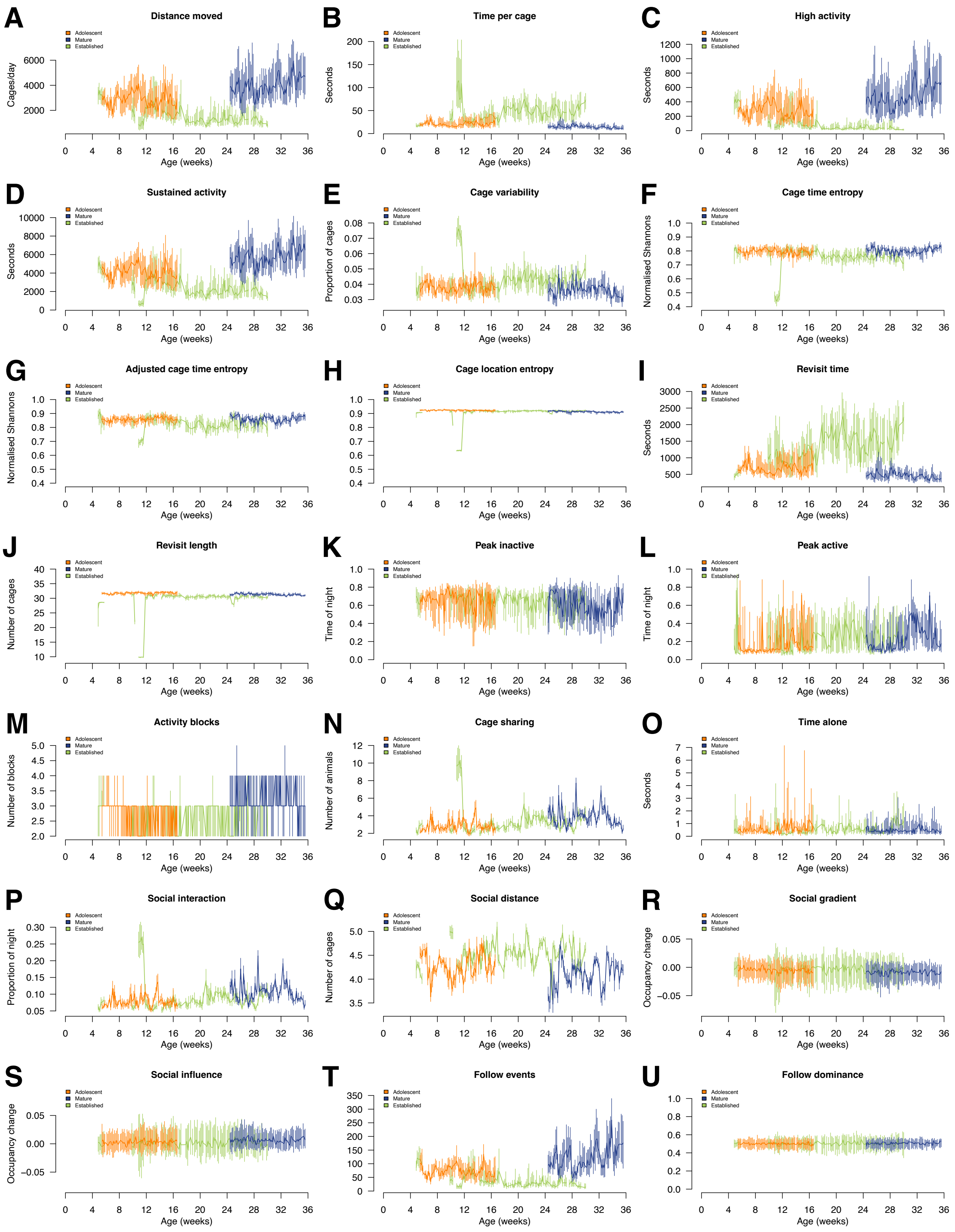
A
Distance moved
Cages/day
Age (weeks)
B
Time per cage
Seconds
C
High activity
D
Sustained activity
E
Cage variability
Proportion of cages
F
Cage time entropy
Normalised Shannons
G
Adjusted cage time entropy
H
Cage location entropy
I
Revisit time
J
Revisit length
Number of cages
K
Peak inactive
Time of night
L
Peak active
M
Activity blocks
Number of blocks
N
Cage sharing
Number of animals
O
Time alone
P
Social interaction
Proportion of night
Q
Social distance
R
Social gradient
Occupancy change
S
Social influence
T
Follow events
U
Follow dominance
Adolescent
Mature
Established

**Fig. S1.**

The 21 metrics calculated by the *ColonyTrack* software present a multimetric description of the behaviour of animals in the home cage tracking environment. Data are presented as group medians for each colony (adolescent: orange; mature: blue; 'established', a colony that had been co-housed since weaning: green) with vertical lines indicating the upper and lower quartiles. Data for all three colonies are plotted together with animal age on the *x*-axis to highlight differences between groups. Some transient anomalies in the cage layout are evidenced by extreme values for certain metrics in the established colony at younger ages.

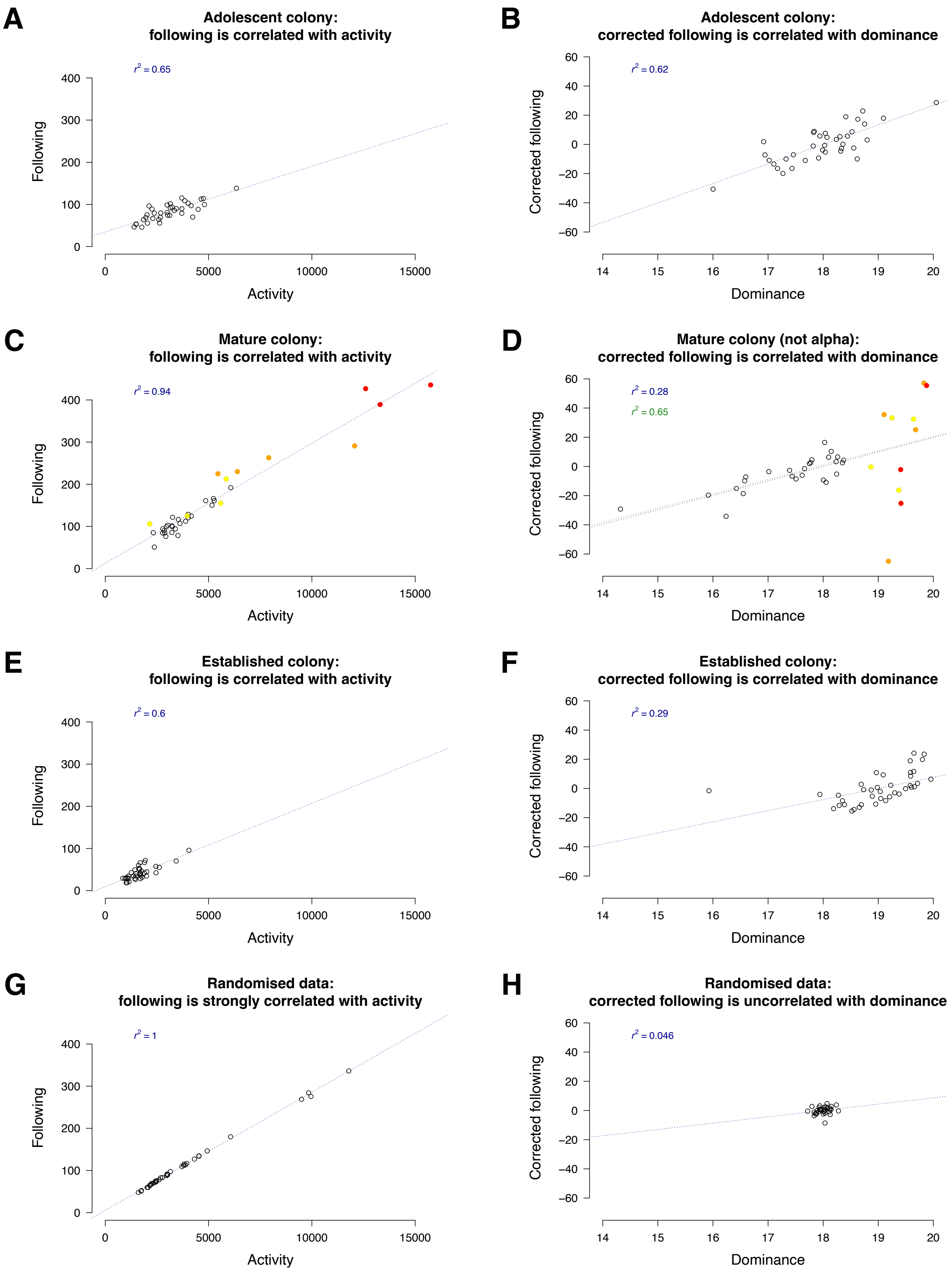
A
Adolescent colony:
following is correlated with activity
$r^2$ = 0.65
Following
Activity
B
Adolescent colony:
corrected following is correlated with dominance
$r^2$ = 0.62
Corrected following
Dominance
C
Mature colony:
following is correlated with activity
$r^2$ = 0.94
Following
Activity
D
Mature colony (not alpha):
corrected following is correlated with dominance
$r^2$ = 0.28
$r^2$ = 0.65
Corrected following
Dominance
E
Established colony:
following is correlated with activity
$r^2$ = 0.6
Following
Activity
F
Established colony:
corrected following is correlated with dominance
$r^2$ = 0.29
Corrected following
Dominance
G
Randomised data:
following is strongly correlated with activity
$r^2$ = 1
Following
Activity
H
Randomised data:
corrected following is uncorrelated with dominance
$r^2$ = 0.046
Corrected following
Dominance

**Fig. S2.**

The activity signal needs to be removed from ‘following’ data to expose the correlation with the social hierarchy. A correlation exists between ‘following’ and activity (‘distance.moved’) in the adolescent (A), mature (C) and established (E) colonies. Residual values from a regression of following against activity are correlated with the dominance (normalised David's scores) for all colonies (B, D, F). The correlation between the activity-corrected following values (which can be thought of as ‘chasing’) and dominance is similar in the mature colony to the other two datasets when the *alpha* group is disregarded. The *alpha* animals are notable in that they do not follow the trend of more corrected following (‘chasing’) with higher dominance. When a background dataset is generated by randomising the mature data, the following metric can be entirely explained by differences in activity and the residual ‘corrected following’ values do not correlate with dominance at all. This indicates that the information content of the following metric is lost when the underlying data are randomised.

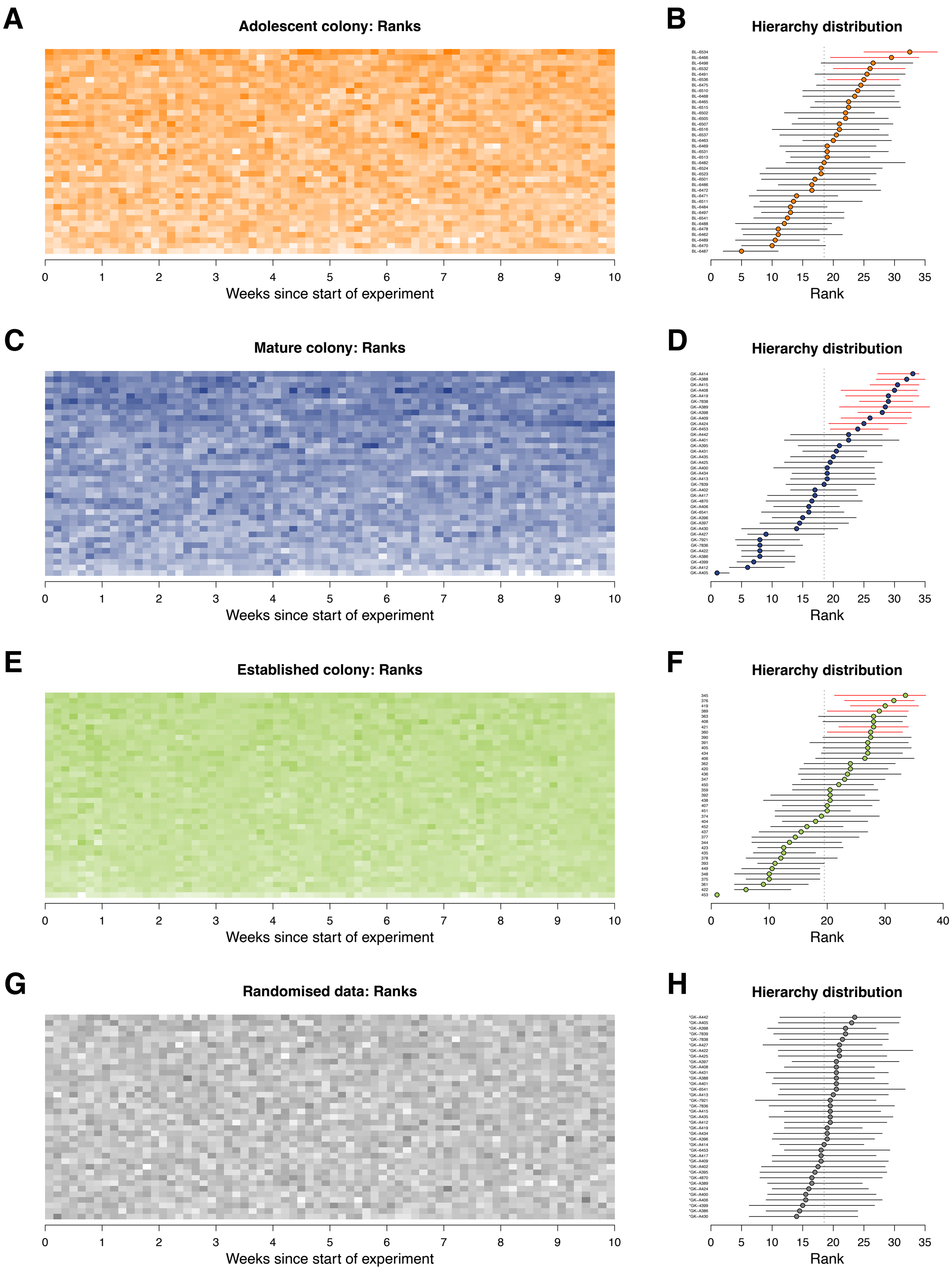
A
Adolescent colony: Ranks
Weeks since start of experiment
B
Hierarchy distribution
Rank
C
Mature colony: Ranks
Weeks since start of experiment
D
Hierarchy distribution
Rank
E
Established colony: Ranks
Weeks since start of experiment
F
Hierarchy distribution
Rank
G
Randomised data: Ranks
Weeks since start of experiment
H
Hierarchy distribution
Rank

**Fig. S3.**

David's scores were calculated and ranked for each day. The heatmaps show darker colours for higher David's score ranks (‘dominance’) for each colony (A, C, E, G). Animals are arranged on the *y*-axis in the same order as for the hierarchy distributions (B, D, F, H). The hierarchy distribution plots for the three tracked colonies (B, D, F) show several high-ranked animals for which the lower quartile of the daily ranks was higher than the median (red horizontal lines). These indicate stability of the ranks and support the assertion that the observed hierarchies are not simply due to chance. Randomising the mature dataset removed the repeatability in dominance ranks so that ranks varied widely over the different days (H).

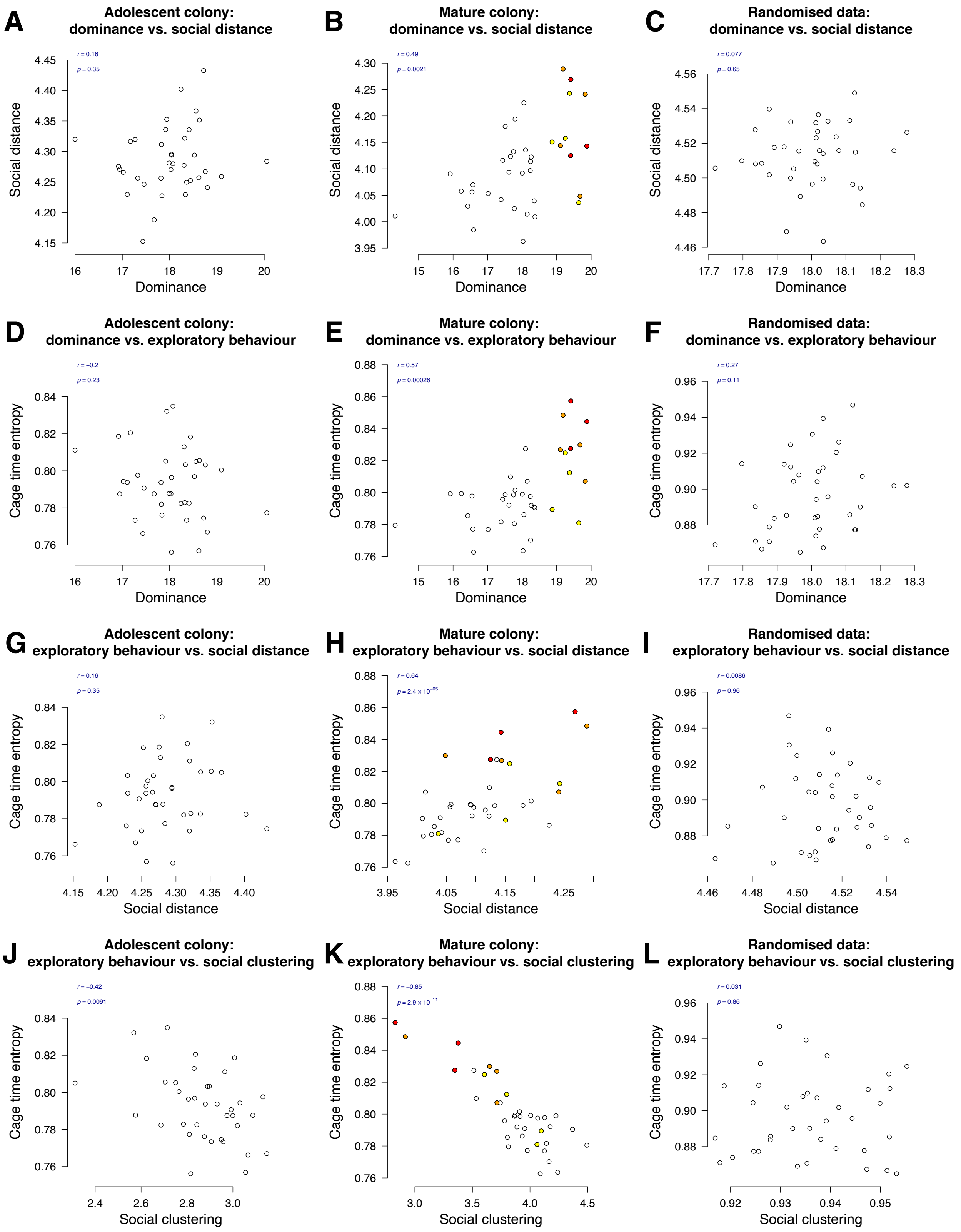
A
Adolescent colony: dominance vs. social distance
r = 0.16
p = 0.35
Social distance
Dominance
B
Mature colony: dominance vs. social distance
r = 0.49
p = 0.0021
Social distance
Dominance
C
Randomised data: dominance vs. social distance
r = 0.077
p = 0.65
Social distance
Dominance
D
Adolescent colony: dominance vs. exploratory behaviour
r = −0.2
p = 0.23
Cage time entropy
Dominance
E
Mature colony: dominance vs. exploratory behaviour
r = 0.57
p = 0.00026
Cage time entropy
Dominance
F
Randomised data: dominance vs. exploratory behaviour
r = 0.27
p = 0.11
Cage time entropy
Dominance
G
Adolescent colony: exploratory behaviour vs. social distance
r = 0.16
p = 0.35
Cage time entropy
Social distance
H
Mature colony: exploratory behaviour vs. social distance
r = 0.64
p = 2.4 × 10−05
Cage time entropy
Social distance
I
Randomised data: exploratory behaviour vs. social distance
r = 0.0086
p = 0.96
Cage time entropy
Social distance
J
Adolescent colony: exploratory behaviour vs. social clustering
r = −0.42
p = 0.0091
Cage time entropy
Social clustering
K
Mature colony: exploratory behaviour vs. social clustering
r = −0.85
p = 2.9 × 10−11
Cage time entropy
Social clustering
L
Randomised data: exploratory behaviour vs. social clustering
r = 0.031
p = 0.86
Cage time entropy
Social clustering

**Fig. S4.**

Social dominance (David's score) was compared against 'social distance' (A–C) and 'cage time entropy', a measure of exploration/cage use, (D–F). The relationship between cage time entropy and social distance (G–I) and 'social clustering' (the average number of animals sharing the same cage; J–L) was also assessed.

Although no relationship was observed in the adolescent colony for any comparison (A, D, G, J), higher dominance in the mature colony was associated with larger social distance (B) and increased exploratory behaviour (E). Exploration, as measured by 'cage time entropy', was also associated with decreased social closeness as evidence by a positive correlation with social distance (H) and negative correlation with social clustering (K). To confirm that these relationships were not statistical artefacts, we repeated each comparison using a randomised version of the data from the mature colony and confirmed the absence of any correlations (C, F, I, L).

### Supplementary Text

Non-agonistic following due to background activity

We had arranged our ColonyRack such that all animals had free access to their sub-network of cages (each experimental group was isolated from the other, but within the groups free social interaction was allowed). This meant that the occasion would often arise, entirely by chance, that two animals would find themselves heading in the same direction and wanting to use the same tunnel simultaneously. In such cases, both animals would pass through the tunnel *in random order* and without conflict. This encounter would nevertheless be picked up as 'following' by the *ColonyTrack* software. We termed this 'neutral' following in **fig. 2D**. Because the following order in such cases is random, this results in a background of 'interactions' with no clear resulting hierarchy information. This emergent feature of the ColonyRack can be demonstrated using randomised datasets (see **fig. S2**). Overlaid on this background are cases of *agonistic* interactions, which are *non-random*. We show that dominant animals tend to be the 'chasers' in such conflicts and this relationship is quite stable (actually surprisingly so, considering the very weak hierarchies present in colonies of isogenic female mice under no resource constraints). Because the evenly-distributed background signal is ignored by the David's score algorithm (due to the winner-takes-all approach in which the results are discretised to a 1 or 0 for net win or net loss respectively), the dominance estimates are only influenced by the directed agonistic interactions.

Inconsistency of out-of-cage behavioral testing versus home cage observations

It has been noted previously (*10*, *11*) that laboratory testing arenas pose an unnatural environment for animals and the results of such tests may not accurately reflect natural behaviour. We observed this in the current experiment where one of the mice performed unexpectedly in the tube test (see **fig. 2E–G**). In this case, the 'omega' animal (the very lowest position in the hierarchy) was seen to lose the majority of its conflicts in the automated ranking from *ColonyTrack* and this status was confirmed by the visual ethology from home cage video footage. However, the same animal exhibited much higher aggression when tested in the tube test, a standard laboratory procedure for assessing dominance in rodents. It is possible that this was due to anxiety induced by the unfamiliar setting.

**Data S1. (separate file)**
Win/loss table from following interactions between CD1 mice in the ColonyRack home cage (file 'WinLoss_CD1CR.tab').

**Data S2. (separate file)**
Ethology of recorded interactions between CD1 mice in the ColonyRack home cage (file 'VideoEthology.tab').

**Data S3. (separate file)**
Experimental dominance data for pairs of CD1 mice over 5 trials in the tube test (file 'CD1_tube_dominance.tab').

**Data S4. (separate file)**
ColonyRack raw data for the adolescent colony as an archived *ColonyTrack* data object (file 'YOUNG.RData' available at https://doi.org/10.5281/zenodo.8333908).

**Data S5. (separate file)**
ColonyRack raw data for the mature colony as an archived *ColonyTrack* data object (file 'OLD.RData' available at https://doi.org/10.5281/zenodo.8333908).

**Data S6. (separate file)**
ColonyRack raw data for the established colony as an archived *ColonyTrack* data object (file 'WIND.RData' available at https://doi.org/10.5281/zenodo.8333908).

**Data S7. (separate file)**
ColonyRack metrics and metadata for the adolescent colony as an archived *ColonyTrack* metrics object (file 'YOUNG_metrics.RData' available at https://doi.org/10.5281/zenodo.8333908).

**Data S8. (separate file)**
ColonyRack metrics and metadata for the mature colony as an archived *ColonyTrack* metrics object (file 'OLD_metrics.RData' available at https://doi.org/10.5281/zenodo.8333908).

**Data S9. (separate file)**
ColonyRack metrics and metadata for the established colony as an archived *ColonyTrack* metrics object (file 'WIND_metrics.RData' available at https://doi.org/10.5281/zenodo.8333908).
